# Strong Components of Epigenetic Memory in Cultured Human Fibroblasts Related to Site of Origin and Donor Age

**DOI:** 10.1101/025288

**Authors:** Nikolay A. Ivanov, Ran Tao, Joshua G. Chenoweth, Anna Brandtjen, Michelle I. Mighdoll, John D. Genova, Ronald D. McKay, Yankai Jia, Daniel R. Weinberger, Joel E. Kleinman, Thomas M. Hyde, Andrew E. Jaffe

## Abstract

Differentiating pluripotent cells from fibroblast progenitors is a potentially transformative tool in personalized medicine. We previously identified relatively greater success culturing dura-derived fibroblasts than scalp-derived fibroblasts from postmortem tissue. We hypothesized that these differences in culture success were related to epigenetic differences between the cultured fibroblasts by sampling location, and therefore generated genome-wide DNA methylation and transcriptome data on 11 intrinsically matched pairs of dural and scalp fibroblasts from donors across the lifespan (infant to 85 years). While these cultured fibroblasts were several generations removed from the primary tissue and morphologically indistinguishable, we found widespread epigenetic differences by sampling location at the single CpG (N=101,989), region (N=697), “block” (N=243), and global spatial scales suggesting a strong epigenetic memory of original fibroblast location. Furthermore, many of these epigenetic differences manifested in the transcriptome, particularly at the region-level. We further identified 7,265 CpGs and 11 regions showing significant epigenetic memory related to the age of the donor, as well as an overall increased epigenetic variability, preferentially in scalp-derived fibroblasts -83% of loci were more variable in scalp, hypothesized to result from cumulative exposure to environmental stimuli in the primary tissue. By integrating publicly available DNA methylation datasets on individual cell populations in blood and brain, we identified significantly increased inter-individual variability in our scalp- and other skin-derived fibroblasts on a similar scale as epigenetic differences between different lineages of blood cells. Lastly, these epigenetic differences did not appear to be driven by somatic mutation – while we identified 64 probable de-novo variants across the 11 subjects, there was no association between mutation burden and age of the donor (p=0.71). These results depict a strong component of epigenetic memory in cell culture from primary tissue, even after several generations of daughter cells, related to cell state and donor age.

## Introduction

DNA methylation (DNAm) at CpG dinucleotides plays an important role in the epigenetic regulation of the human genome, contributing to diverse cellular phenotypes from the same underlying genetic sequence. For example, DNAm levels at particular genomic loci can accurately classify different tissues (1) and even underlying cell types within tissues (2). These stable cell type- and tissue-discriminating loci appear to represent only a subset of “dynamic” CpGs, approximately 21.8%, actively involved in regulation of gene expression (3). Changes in these epigenetic patterns across aging have been extensively studied (4), particularly in large studies of whole blood (5–7), but subsets of these age-associated CpGs appear tissue-independent (8).

These epigenetic barcodes also play an important role in cellular reprogramming (the conversion of somatic cells to pluripotent stem cells), a powerful and promising experimental system in biology, genetics and personalized medicine (9). This epigenetic reprogramming of somatic cells to induced pluripotent stem cells (iPSCs) induces demethylation (10) followed by specific patterns of subsequent DNA methylation that can reflect the original somatic tissue (11). Fibroblasts are one of the most popular cell types for generating iPSCs (12), particularly from skin, given the relative ease of access to these cells, although other skin-derived cell types such as keratinocytes from the same individual generate similar iPSC cell lines (13). Skin, however, is perhaps the most susceptible tissue source in the body to environmentally induced insult, particularly through sunlight and chemical exposures, which can induce changes in epigenetic patterns (14). The epigenetic “memory” of source tissue for iPSC characterization has been well characterized (11).

In our previous work, we successfully cultured fibroblast lines from the dura mater of postmortem human donors, a source location largely protected from environmental insult with slowly dividing cells (15). We compared these cultured fibroblast lines to those derived from scalp samples from the same individuals, and found that the rate of culture success was higher for dura-derived fibroblasts; in some cases only the dura fibroblasts from an individual would culture. While the resulting cultured cells from these two sampling locations were largely morphologically indistinguishable (see Figure 1 in Bliss et al, 2012), we hypothesized that increased culture success might have a strong epigenetic component. Previous reports have indicated that cultured cells have largely stable epigenomes, with the exception of a small number of loci (16). We therefore sought to characterize the methylomes and transcriptomes of fibroblasts from these two sampling locations – scalp and dura – from donors across the lifespan.

**Fig. 1.**
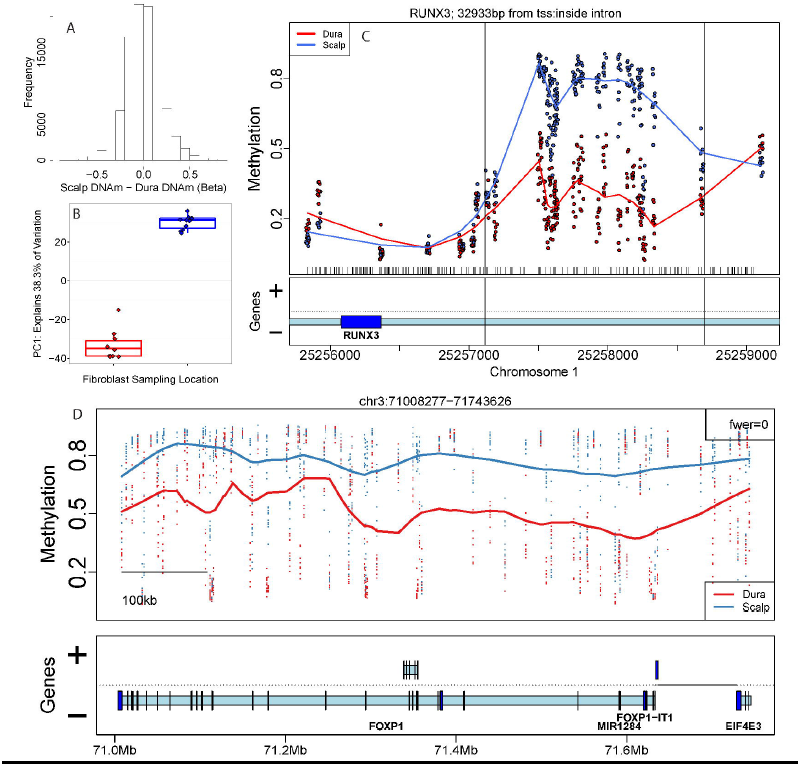
DNA methylation patterns in dura- and scalp-derived fibroblasts. (A) Histogram of difference in DNAm levels at CpGs/probes between scalp and dura derived fibroblasts (on the proportion methylation scale). (B) The first principal component (PC1) of the DNAm data plotted against fibroblast sampling location (scalp versus dura). (C) Example significant differentially methylated region (DMR) that overlaps the gene *RUNX3*, with DNAm levels on the y-axis and genomic coordinates on the x-axis. (D) Example significant DNAm block, with DNAm levels on the y-axis and genomic coordinates on the x-axis. Gene annotation panels in C and D are based on Ensembl annotation – dark blue represents exons and light blue represents introns.

Here we identify several components of epigenetic “memory” in cultured fibroblasts after multiple passages (i.e. splitting and continuing to grow) where primary tissue originated from two locations in the body. The strongest epigenetic memory was related to sampling location in the body, as we identified widespread DNAm differences at local and regional spatial scales preserved through identical culturing processes. We further find increased stochastic epigenetic variability in cultured fibroblasts from the scalp compared to dura. This increased variability manifested in significant increased quantitative pairwise epigenome-wide distances in a combined analysis with publicly available DNAm data on skin fibroblasts (17), pure cell populations from peripheral blood (18), and cells from the dorsolateral prefrontal cortex (19). Another component of epigenetic memory was related to the age of the donor, including a subset of CpGs that displayed location-dependent changes through aging. The epigenetic differences between these fibroblasts appear to occur largely through epigenetic-dependent mechanisms, as there were few differences in coding sequence across the fibroblasts from the two locations within the same individual. These results demonstrate the effect of epigenetic memory in cultured fibroblasts by sampling location and donor age in morphologically indistinguishable cells.

## Results

We measured DNA methylation (DNAm) levels from scalp- and dura-derived cultured fibroblasts in 11 postmortem donors (22 samples) from across the lifespan, ranging from early infancy to 85 years (S1 Fig., S1 Table), using the Illumina HumanMethylation450 microarray (Illumina 450k) (20). After data processing, normalization, and quality control with the minfi package (21), we obtained normalized data on 21 samples (one dura sample with lower quality was removed prior to across-sample normalization) across 456,513 probes (probes with single nucleotide polymorphisms, SNPs, at the target CpG or single base extension site were removed, as were probes on the sex chromosomes, see Methods).

### Strong components of epigenetic memory by primary cell sampling location

We first characterized differences in DNAm levels from cultured fibroblasts derived from different locations (scalp versus dura). Many probes, targeting individual CpGs, were differentially methylated between scalp- and dura-derived fibroblasts – 101,989 (22%) at genome-wide significance (false discovery rate, FDR < 5%, see Methods). These significant DNAm differences between cultured fibroblasts from the scalp and dura were large in magnitude, with 57,704 probes having differences in DNAm levels greater than 10%, and 23,752 with differences greater than 20% (Fig. 1A). The directionality of these DNAm differences were balanced, with approximately equal proportions of CpGs showing increased versus decreased methylation in cultured fibroblasts from scalp compared to dura. These differentially methylated probes (DMPs) were widely distributed across the genome, as 18,551 genes (defined by UCSC knownGene database, see Methods) had at least one DMP within 5 kilobases (kb). These widespread single CpG differences manifest as the largest component of variability in the entire dataset, as the first principal component (Fig. 1B, explaining 38% and 62.3% of the variability before and after surrogate variable analysis, SVA (22)) represents the sampling location of these cultured fibroblasts, suggesting a strong epigenetic memory of original cell location.

Since these differentially methylated CpGs tended to cluster in a smaller number of genes, we further identified 697 differentially methylated regions (DMRs) at stringent genome-wide significance (family-wise error rate, FWER < 10%) – these regions were identified based on adjacent probes showing directionally-consistent differences in DNAm > 10% between groups (23) (see Methods). For example, we identified a region of 24 contiguous probes hypermethylated in scalp-derived fibroblasts within the gene *RUNX3* – a tumor suppressor that plays an integral role in regulating cell proliferation and the rate of apoptosis (24) (Fig. 1C, see S2 Fig. and S2 Table for all significant DMRs). Regional differences, particularly in CpG island shores, previously have been shown to better distinguish tissues and cell types (1) and correlate with neighboring gene expression levels (21) than individual CpGs. Unlike at the single CpG level, which had balanced directionality of differential methylation, the majority of DMRs had higher DNAm levels in fibroblasts derived from scalp compared to those derived from dura (N=414, 59.4%). Using gene sets defined by biological processes (25), these neighboring genes (within 5 kb) were strongly enriched for morphogenesis (including morphogenesis of the epithelium), developmental processes, cell differentiation, and epithelium and connective tissue development, among other more general gene sets (all p < 10^−8^, S3 Table).

In addition to the extensive differential methylation at both the CpG and regional level, we identified 243 long-range regions with consistent significant methylation change (FWER < 10%), called “blocks” (26), using an algorithm adapted from whole genome bisulfite sequencing (WGBS) data to Illumina 450k (21). A representative significant block is shown in Fig. 1D (see S3 Fig. for all significant blocks at FWER < 10%). Blocks have now been identified across many cancer types (27), and tend to associate with higher order chromatin structure including nuclear lamin-associated domains (LADs) (28) and large organized chromatin K9 modification (LOCKs) (26). The 243 significant blocks in our data represent 41 Mb of sequence and contain 298 annotated genes. These blocks contain 41 of the significant DMRs that differentiate sampling location of the fibroblasts, and more interestingly, every block overlaps at least one “dynamic” cell/tissue DMR identified using WGBS data from Ziller et al (2013) (3).

While these cultured fibroblasts were several generations/passages removed from the primary tissue and morphologically indistinguishable, we nevertheless found widespread epigenetic differences by sampling location of the primary fibroblasts at varying spatial scales, suggesting a strong epigenetic memory of the original cell location.

### Epigenetic memory related to original cell location manifests in the transcriptome

We next sought to determine the functional correlates of the widespread epigenetic differences identified between scalp- and dura-derived fibroblasts by performing RNA sequencing (RNA-seq) on polyadenylated (polyA+) m RNA from the same cultured samples (see Methods). Briefly, we aligned the reads to the transcriptome using TopHat (29) and generated normalized gene counts (as fragments per kb per million mapped reads, FPKM) based on the Illumina iGenome hg19 annotation using the featureCounts software (30). A median of 88.0% (interquartile range, IQR: 85.5% – 88.8%) of reads mapped to the genome, of which a median of 84.7% (IQR: 84.4%-85.5%) mapped to the annotated transcriptome (see S1 Table for sample-specific percentages). We identified 11,218 expressed genes with average FPKM expression greater than 1.0. Initial clustering of the FPKM values separated the fibroblast samples by location in the first principal component (PC), which explained 35.4% of the variance (S4 Fig.), mirroring the first principal component of the DNAm data (Fig. 1B). Differential expression analysis of the RNA-seq data, independent of the results from the epigenetic analyses above, identified many genes that differed by the source of the primary fibroblast – 5,830 genes at FDR < 5%. These genes were strongly enriched for signaling and cell communication, cell proliferation, apoptotic processes, and epithelium development and morphogenesis via gene ontology (GO) analysis (all p < 10^−8^, S4 Table).

We next used the gene expression data as a functional readout of the differentially methylated loci identified between fibroblasts cultured from scalp and dura. The majority of significant DMPs (76,971/101,989, 75.47%) were inside or near (within 5kb of) a UCSC annotated gene, and 28.2% (21,742/76,971) were significantly associated with gene expression levels (at p < 0.05). This percentage of DMPs with significant expression readout was elevated (34.9%) among those DMPs with larger DNAm differences by sampling location (greater than 10% difference in DNAm levels). These DMPs were strongly significantly enriched among the CpG sites that associated with expression levels at the p < 0.05 (48,062 probes within 5kb of genes, odds ratio, OR=3.99, p < 2.2×10^−16^) and FDR < 0.05 (6,559 probes within 5kb of genes, OR=19.54, p < 2.2×10^−16^) significance thresholds.

Similarly, 587/697 (84.2%) DMRs were in or near (<5kb) genes, and many had DNAm levels that were significantly associated with gene expression levels (306/587, 52.1% at p < 0.05). For instance, a DMR overlapping an intronic sequence of the *SIM1* gene (Fig. 2A) was unmethylated with low corresponding expression of the gene in the cultured fibroblasts from dura, and highly methylated with corresponding high expression levels of the gene in the scalp-derived fibroblasts (Fig. 2B and S2 Table). This is in line with previous reports suggesting that gene body methylation levels positively associate with local gene expression (31), unlike CpG island shore methylation that tends to be negatively associated with gene expression levels (1). Of the 478 unique genes in or within 5kb of DMRs, the expression of 235 (49.2%) of them was significantly correlated with DNAm (p < 0.05). These 235 unique genes tended to exhibit stronger differential expression between the scalp- and dura-derived fibroblasts (median fold change = 1.59, IQR = 1.23-2.68) than individual CpG results, in line with previously published findings (21). GO analysis on expression-associated genes proximal to DMRs revealed enrichment for multiple important biological processes such as connective tissue development, epithelium morphogenesis and development, cell differentiation (specifically including epithelial cell differentiation), and cell proliferation (specifically including epithelial cell proliferation), among other more general sets (all p < 10^−8^, see S5 Table).

**Fig. 2.**
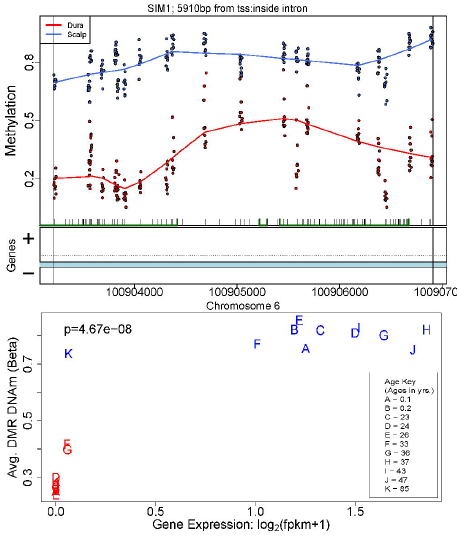
Regional DNA methylation changes manifest in the transcriptome. (A) Plot of the DNAm levels (proportion methylation) of an example significant DMR, which overlaps the gene SIM1. (B) Plot of the average DMR DNAm levels versus the expression level of *SIM1*, showing high positive correlation (p=4.67×10^−8^).

Lastly, we found that the majority of differentially methylated blocks contained at least one gene differentially expressed between scalp- and dura-derived fibroblasts. While the majority of blocks contained at least one gene (N=188/243, 77.4%), 63.8% (N=120/188) had at least one gene that was differentially expressed (at p < 0.05). For example, one of the blocks, hypermethylated in scalp-derived fibroblasts, overlaps the *HOXB* gene cluster (Fig. 3A). In this block, expression levels of the *HOXB* genes are significantly greater in fibroblasts cultured from scalp than those from dura (Fig. 3B). Similarly, the 188 significant blocks contained 298 unique genes, and 126 of them (42.3%) were differentially expressed (at FDR < 0.05) which is a higher proportion than the rest of the transcriptome (0.42 vs. 0.32, p=3.79×10^−9^). These results suggest that epigenetic memory related to original cell location do largely read out in the transcriptome, particularly among regional changes in DNAm related to fibroblast sampling location.

**Fig. 3.**
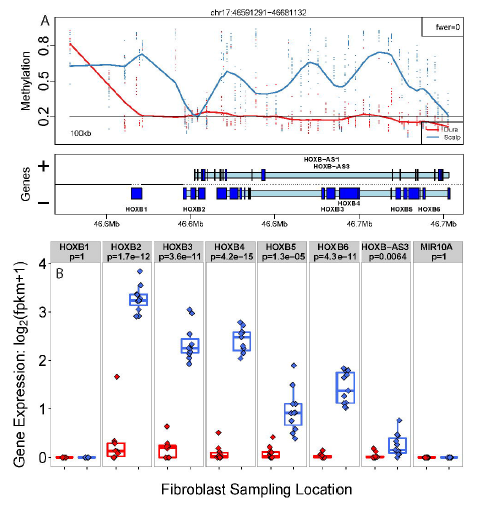
Long-range DNA methylation changes manifest in the transcriptome. (A) Plot of the DNAm levels (proportion methylation) of a significant DNAm block overlapping genes in the *HOX* family. Y-axis: proportion DNAm levels, x-axis: genomic coordinates on chromosome 17. (B) Corresponding expression levels of the *HOX* genes within the DNAm block are more highly expressed in the scalp. Y-axis: log_2_ transformed fragments per kilobase per million mapped (FPKM), x-axis: sampling location.

### Increased stochastic variability in scalp-derived fibroblasts

We hypothesized that scalp-derived fibroblasts might have more variable levels of DNAm than dura-derived fibroblasts, given the chronic exposure to environmental factors (e.g. sunlight, chemicals) in the primary tissue across the lifespan. At the individual CpG level, we tested for differences in variance between the scalp- and dura-derived fibroblasts independent of the underlying mean methylation levels (32) (see Methods section). While only two probes reached genome-wide significance (at FDR < 0.05) for differences in variance, at marginal levels of significance (p < 0.05), fibroblasts cultured from scalp had more variable DNAm levels than fibroblasts cultured from dura (N=13,169/16,330, 80.6%).

We next sought to characterize epigenome-wide patterns of DNAm across these fibroblasts in the context of other diverse cell types. After downloading and normalizing Illumina 450k data from sorted blood (18) and frontal cortex (19), as well as skin-derived fibroblasts (17) and melanoma samples (SKCM) from the Cancer Genome Atlas (TCGA) (33), we computed epigenome-wide Euclidean distances between and across each of the 11 cell types (see Methods section). We noted that these cell types largely cluster by tissue source (brain, blood, and fibroblasts in the first two principal components and largest dendrogram splits, S5 Fig.).

The inter-individual epigenomic distances, and their variability, were much greater in the scalp-derived (as well as skin-derived) fibroblasts than dura-derived fibroblasts (p=1.34×10^−9^ and p=1.77×10^−14^ respectively, see Fig. 4). The distances within scalp- and skin-derived fibroblasts were significantly larger than those calculated within pure blood and cortex cell types (p-values range from 1.04×10^−21^ to <10^−100^). Interestingly, the inter-individual distances between fibroblasts cultured from scalp samples were greater than the distances between different cell types within a blood cell lineage (e.g. natural killer cells versus CD4+ T-cells) and similar to distances across lineage (e.g. natural kill cells versus monocytes). Note that comparing inter-individual distance between two cell types (e.g. scalp-versus dura-derived fibroblasts) reflects the extensive differential methylation between these two cell types (see Fig. 1) – the inter-individual distances are large but the variability in distances was low.

**Fig. 4.**
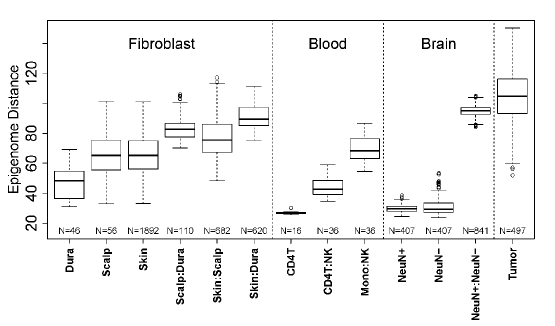
Increased epigenome distances within scalp-derived fibroblasts. Y-axis: epigenome (Euclidean) distance between pairs of samples stratified by cell and tissue types. CD4T: CD4+ T-cell, NK: natural killer cell, Mono: monocyte, NeuN+: neuronal DLPC cell, NeuN-: non-neuronal DLPFC cell.

As another example, the distances across scalp-derived fibroblasts were less than comparing inter-individual variability between neurons and non-neurons (via NeuN+ sorting), which reflects the extensive methylation differences between these two cell types (19). As expected, we found the greatest epigenome-wide distances and largest inter-individual variability in the melanoma samples (26, 32), which highlights the relative scale of these epigenome-wide distances (ranging from pure cell types to cancer). These increased epigenomic distances may relate to the rate of cell division, which is non-existent in neuronal cells (34) and infrequent in T-lymphocytes at the population level (35). The increased epigenetic variability in the scalp samples was further not associated with differences in donor age (p > 0.05, S6 Fig.), suggesting increased epigenetic stochastic variability in scalp- (and skin-) derived fibroblasts.

### Epigenetic memory related to donor age

We hypothesized that a subset of this increased variability might result from age-related divergence in DNAm at individual loci that were differential by sampling location, such that young donors would have lesser difference in DNAm levels, and older donors would have larger differences in DNAm. By fitting linear models on the difference in DNAm levels across sampling location as a function of donor age (see Methods), we identified 7,265 CpGs associated with diverging DNAm levels across aging (at FDR < 10%, S7 Fig.). These loci appeared to be clustered into representative patterns of their age-related changes (Fig. 5). The majority of these CpGs had significant age-related changes in fibroblasts derived from the scalp (64.0%), but not dura (17.4%), and the magnitude of change across age was larger in scalp-derived fibroblasts – the average change in percent DNAm per decade of life was 3.13% (IQR = 1.81%-4.29%) in fibroblasts derived from scalp compared to 1.13% (IQR = 0.295%−1.61%) in those from the dura mater.

**Fig. 5.**
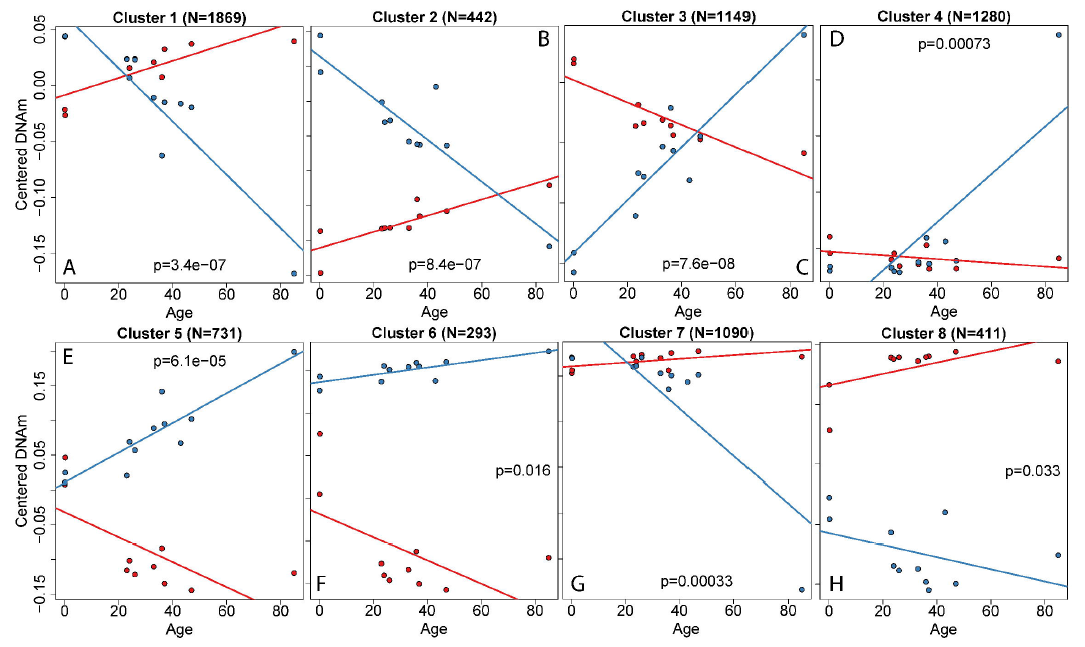
Representative patterns of age-associated changes in DNAm by sampling location. Y-axis: mean-centered DNAm levels, y-axis: sample age, p-value represents the statistical interaction between sampling location and age on DNAm levels. N: number of CpGs in the cluster.

A subset of these CpGs showing sampling location-dependent age-related changes associated with nearby gene expression levels. Most of the probes (N=5,185/7,265, 71.4%) were annotated to 3,553 unique genes (within 5kb) and 21.8% of these (N=775/3,555) showed significant correlation between DNAm and gene expression (p < 0.05). These DNAm associated genes were enriched for multiple general developmental processes including cell development, morphogenesis, and differentiation (all p<10^−8^, S6 Table). Several of the age-related CpGs showing expression association were within genes that are involved in cell proliferation and apoptosis. For instance, DNAm levels at two significant probes inside the gene *TEAD1*, which regulates notochord development and cell proliferation (36), were significantly associated with gene expression levels (p=8.60×10^−4^ and 0.045, respectively). Another significant DNAm-expression pair (p=0.02) was located in *AVEN*, a gene shown to inhibit Caspase activation in apoptosis (37). These results suggest altered regulation of DNAm levels across aging occurs primarily in fibroblasts derived from scalp but not from dura, perhaps through altered cell proliferation and apoptosis, and possibly reflecting greater exposure to environmental agents that can affect the epigenome.

### Epigenetic memory related to sampling location and age do not implicate genetic mosaicism

Lastly, we characterized the expressed sequences of the scalp- and dura-derived fibroblasts within each individual to examine the extent of genetic mosaicism, which may contribute to differences in DNAm through changing the underlying genetic sequence in the fibroblasts taken from scalp. *De novo* variants were called directly from the RNAseq data, and after filtering by many quality metrics (see Methods) we identified 64 high-confidence candidate variants that were discordant by sampling location in at least a single individual (S7 Table), including 22 annotated coding variants (13 synonymous and 9 non-synonymous) (38). We found no association between coding variant burden and subject age (p=0.71, S8 Fig.). These results suggest that many of the location- and age-associated DNAm differences are not due to somatic mosaicism and likely arise through epigenetic mechanisms that are maintained through cell culture and multiple passages.

## Discussion

Here we interrogated the epigenomes and transcriptomes of pairs of fibroblasts cultured from scalp and dura mater taken from the same individual, in a subject cohort that ranges in age across the human lifespan. These cultured fibroblasts, generations removed from the primary tissue of origin, and with indistinguishable morphology, still maintained strong components of epigenetic “memory” related to sampling location (scalp versus dura) and differential changes in DNAm levels across aging. The widespread differences in DNAm levels by sampling location were identified at many spatial scales, including single CpGs, differentially methylated regions, blocks, and globally. Furthermore, many of these differences in DNAm levels manifested in the transcriptome, showing significant corresponding differences in expression for genes most proximal to these epigenetic changes. The genes with differences in expression and DNAm levels by sampling location were previously implicated in processes relating to cell proliferation and apoptosis, which likely relate to the function of the fibroblasts in the primary tissue. One might have predicted this outcome, as fibroblasts in the scalp, including those that are cultured turnover much more rapidly than those in the dura mater (15).

Another component of epigenetic memory in these cultured fibroblasts was related to ages of the donors, where age-related changes occurred differentially by sampling location. These age-associated loci can be clustered into general patterns of epigenetic changes by age and location, all showing significant interaction between donor age and sampling location. While some patterns were expected, such as divergence in DNAm levels from similar levels at birth (clusters 1, 4, 5, and 7), several other clusters showed an unexpected convergence in DNAm across aging (clusters 2 and 3). We do note that the elderly donor (age 85) is influential in both the statistical discovery at individual loci and in some of the subsequent clusters – larger sample sizes can hopefully further define and replicate these observations. Also, while the fibroblasts were analyzed from some subjects with psychiatric disorders, almost all comparisons between scalp and dura sampling locations, and differential changes with age were naturally matched within an individual, reducing the potential impact of diagnostic confounding. Furthermore, a larger sample size would likely identify significant divergence in DNAm at the region level – while we found 7,265 individual CpGs, we found very few DMRs at global significance (6 and 11 DMRs at FWER ≤ 10% and 20% respectively). The region-finding approach has been shown to be statistically conservative (23) and the identification of these differential age-related changes by sampling location was based on number of donors (N=10), not the number of observations (N=21).

These age-related changes in cultured fibroblasts are one of the first examples, to our knowledge, of genome-wide significant age-related changes in a pure cell population that is many mitoses and passages from the original donor cells. Many papers have identified widespread age-related changes in heterogeneous cell populations, like blood (5, 7), brain (39), and other tissue types (8), which may result in false positives when the underlying cellular composition changes across aging (4). Other papers have used individual cell populations to validate age-associated loci identified in homogenate tissue at marginal significance (40) or have identified age-related changes in targeted approaches at limited number of loci (41).

Similarly, these fibroblasts cultured from the scalp and dura mater were the first example, again to our knowledge, of morphologically indistinguishable cells with vastly different epigenomic profiles. Using epigenomic distances, these two cohorts of fibroblasts were more different in their DNAm patterns than different lineages of blood cells, while less different that neuronal versus non-neuronal cells from the frontal cortex (Fig. 4); the cells underlying each comparison have very different morphologies and cellular function. Furthermore, the majority of differences in DNAm levels between scalp- and dura-derived cultured fibroblasts appeared to be determined early in development, prior to early infancy in this sample, and remained stable throughout the lifespan. Of the 101,989 significant DMPs for sampling location, 98,461 (96.5%) were not associated with differential age-related changes. These findings demonstrate strong components of epigenetic memory related to cell location and aging in fibroblasts cultured from the scalp and dura mater from postmortem human donors.

There are important implications from this study for the field of regenerative medicine. If fibroblasts are going to be the source for iPSCs, and ultimately differentiated tissues, the source of these fibroblasts, and their epigenetic characteristics, may be an important consideration. For example, these differences in cellular states in cultured fibroblasts may relate to the number of cell divisions, as skin and scalp fibroblasts have a much quicker turnover than fibroblasts in the dura (15). The extent of cell division could relate to the epigenomic distances between and across the diverse cell types we have analyzed. Further research may better determine the extent of epigenetic memory of cell state of fibroblasts cultured from different locations after the generation of iPSCs and subsequent differentiation into new cell types. As the field of regenerative medicine advances, our study demonstrates that deciding upon the source of fibroblasts from an individual to generate new tissues and organs may be an important consideration. While it was shown that transcriptional variability by tissue of origin was low in iPSCs [13], it was also demonstrated that the DNAm landscape in iPSCs differs greatly by tissue or origin, and this phenomenon may explain the propensity of iPSCs derived from different somatic tissues to differentiate into different lineages [11].

## Methods and Materials

### Human Tissue Collection

Human dural and scalp fibroblasts on which the methylation and gene expression studies were performed were obtained from fibroblast cell lines derived from human post mortem scalp and dura mater tissues. For this study, tissues from 11 individuals were used, with the ages of individuals ranging from 0.1 to 85 years of age (see S1 Table for additional demographics). The postmortem tissues from 2 of the subjects were collected by the Lieber Institute for Brain Development (LIBD) and the tissues from the remaining 9 subjects were collected by National Institute for Mental Health (NIMH) (Clinical Brain Disorders Branch (CBDB), Division of Intramural Research Programs (DIRP)). The NIMH tissues were collected from two medical examiners (Washington, DC office and Commonwealth of Virginia, Northern District office); the LIBD tissues were obtained the Office of the Chief Medical Examiner (Baltimore, MD). A preliminary neurological or psychiatric diagnosis was given to each case after demographic, medical, and clinical histories were gathered via a telephone screening on the day of donation. For each case, the postmortem interval (PMI) (the time (in hours) elapsed between death and tissue freezing) was recorded. (See Tbl. 1 for PMIs and demographics for every subject used in this study). Every case underwent neuropathological examinations to screen for neurological pathology. Additionally, the medical examiner’s office performed toxicology analysis of every subject’s blood to screen for drugs.

Dura and scalp tissue were collected at the time of autopsy. From the autopsy room, the tissues were transported in separate bags: one containing cerebral dura mater and the other a 1 in × 1 in scalp segment with hair attached. Both bags were transported on wet ice to the lab, where the culture procedure was immediately started.

### Scalp and Dura Tissue Cultures

The dura culture medium was prepared out of 1× DMEM (Ref#11960-044, GIBCO) with 10% by volume fetal bovine serum, 2% by volume 100X GlutaMAX (Cat#: 35050, GIBCO), 1% by volume Penicillin-Streptomycin/Amphotercin solution (Ref# 15140-122, GIBCO), and 1% by volume Gentamicin solution (Cat# 17105-041, Quality Biological). This culture medium was used in all subsequent steps of the dura culturing procedure. The scalp culture medium used for all subsequent steps of the scalp culturing procedure was made the same way except without the 1% Gentamycin. A rinsing solution was prepared out of 1X PBS (pH 7.2) (Ref# 21-040-CV, Corning Life Sciences), 1% by volume Penicillin-Streptomycin/Amphotericin solution (Ref# 15140-122, GIBCO), and 1% by volume Gentamicin (Cat# 17105-041, Quality Biological).

The dissected scalp sample was washed with the rinsing solution three times, the fat tissues were cut away, and all hair was plucked out with forceps. The scalp sample was then placed epidermis side down on a dish and floated with Dispase II enzyme solution (2.4 units of the Dispase II enzyme per mL of PBS, Dispase II enzyme: Cat#17105-041, GIBCO). (Dispase II enzyme is a proteolytic enzyme used to separate the dermis from the epidermis by cleaving the zone of the basement membrane.) The dish was covered with parafilm and foil, and placed in a 37°C incubator for 24 hours. After the 24-hour period, the epidermis was peeled away from the dermis. The dermis was washed with the rinsing solution, dried, and cut into 2-3 mm^2^ pieces. The pieces were placed in a Falcon Easy Grip tissue culture 35×10 mm dish and one drop of scalp culture medium was added to each piece of scalp. The dish was placed in the incubator at 37°C and 5% CO_2_ for culturing.

A similar procedure was followed for the dura samples. Dura samples were washed with the rinsing solution three times. Then, a few 2–3 mm^2^ pieces were cut from the dura mater and placed together in an Easy Grip cell culture 35×10 mm dish. One drop of dura culture medium was added to each dura piece. The culture dish was then placed in an incubator (at 37°C and 5% CO_2_) for culturing. The medium of each culture was changed to fresh medium 2–3 times per week to promote growth of the fibroblasts. On average, fibroblast cells started to proliferate at 7–14 days, however some samples took longer (up to 3 weeks).

### Fibroblast Cell Cultures

The dura and scalp tissue cultures were monitored under a phase-contrast microscope. When the fibroblast growth reached 90-95% confluence, 1 mL of a 0.25% trypsin solution (Cat#T4049, Sigma) was added to each culture dish, and the cells were incubated for 5 to 8 min. Then, 1mL of media was added to each dish stop the enzymatic reaction. Next, the contents of each culture dish were transferred into separate 15 mL Falcon conical tubes and 8mL of media was added to each tube. The conical tubes were centrifuged for 5 min at 1100 rpm. The supernatant was discarded, 5mL of fresh media was added to each conical tube, and the contents of the tubes were transferred onto separate 25 cm^3^ cell culture Easy Flasks (Thermo Scientific, Cat# 156367), where they were kept in cultures for 3-5 days in an incubator (at 37°C and 5% CO2). When the cells reached 90-95% confluence, the cells from each 25 cm^3^ flask were transferred onto two 75 cm^3^ cell culture easy flasks (Thermo Scientific, Cat# 156499) and kept in cultures for continued growth. When the cells reached 90-95% confluence, they were incubated with 3 mL of 0.25% trypsin solution for 5 to 8 min, after which 3mL of fresh culture media was added to stop the enzymatic reaction. Then, the contents of the flasks were transferred into separate 15 mL Falcon conical tube and 4mL of media was added to each tube. The tubes were centrifuged (5 min, 1100 rpm), the supernatant was discarded and the pellets containing the fibroblasts were removed from the centrifuge tubes and transferred to cryoTube vials (Cat#375418, Thermo Scientific). 0.5 mL of recovery cell culture freezing medium (Cat#12648-010, GIBCO) was added to each vial, after which the vials were insulated with Styrofoam and placed into a −80°C freezer. Later, the tubes were transferred to a −152°C liquid nitrogen freezer.

These frozen dura and scalp fibroblast cells were then used generate DNA methylation and gene expression levels.

### DNA Methylation Data Generation

DNA methylation landscapes of the dura- and scalp-derived fibroblasts were analyzed using the Illumina HumanMethylation 450 BeadChip array (“450k”). The 450k array interrogates >485,000 DNA methylation sites (probes) and measures the proportion DNA methylation at each target site (the 450k array interrogates both CpG and CH sites). The microarray preparation and scanning were performed in accordance with the manufacturer’s protocols. The resulting data from the 450k consists of R(ed) and G(reen) intensities using two different probe chemistries (20), which we converted to M(ethylated) and U(nmethylated) intensities using the *minfi* Bioconductor package (21). One dura sample had lower median probe intensities and was removed prior to normalization and downstream analyses. After quality control (QC), the M and U intensities were normalized separately across samples using stratified quantile normalization (21). Probes containing common SNPs (based on dbSNP 142) at the target CpG or single base extension site, and probes on the sex chromosomes were removed, leaving 456,513 probes on 21 samples for analysis.

### Differential methylation analysis

We determined differential methylation using linear modeling on the normalized DNAm levels, using the model:

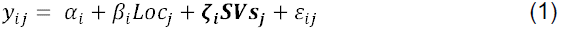

where *y*_*ij*_ is the normalized proportion methylation at probe *i* and sample;*j*, *α*_*i*_ is the proportion methylation in the fibroblasts sampled from the dura mater, *β*_*j*_ is the difference in methylation in the scalp-derived fibroblast, and *Loc*_*j*_ is the sampling location represented by a binary variable (Dura=0, Scalp=1). These statistical models were adjusted for surrogate variables (6 SVs) estimated using surrogate variable analysis (SVA) (22).

Differentially methylation probes (DMPs) were identified by fitting Eq. 1 to each probe, and obtaining the corresponding moderate t-statistic and p-value using the limma package (42). P-values were adjusted for multiple testing using the false discovery rate (FDR) (43) and significant probes were called were FDR < 0.05. Principal component analysis (PCA) was performed after regressing out the surrogate variables from the DNAm levels of each probe, preserving the effect of fibroblast sampling location. Finding differentially methylated regions (DMRs) involves identifying contiguous probes where *β* ≠ 0 using the *bumphunter* Bioconductor package (version 1.6.0) (23), here requiring |*β*| > 0.1 (argument: cutoff=0.1) and assessing statistical significance using linear modeling bootstrapping with 1000 iterations (argument: nullMethod=bootstrap’ and B=1000). DMRs were called statistically significant when the family wise error rate (FWER) ≤ 0.1. We identified blocks using the same model as above using the *blockFinder* function in the minfi package (21), which collapses nearby CpGs into a single measurement per sample, and then fits Eq.1 above, only here *j* represents probe group, not probe. Here we again required at least a 10% change in DNAm between groups and assessed statistical significance using the FWER based on 1000 iterations of the linear model bootstrap.

### RNA extraction and sequencing

RNA was extracted from the cultured dura and scalp fibroblasts with the RNeasy kit (Qiagen), in accordance with the manufacturer’s protocol. RNA molecules were treated with DNase, polyadenylated (polyA+) RNA was isolated, and resulting sequencing libraries were constructed using the Illumina TruSeq RNA Sample Preparation Kit (v2) and sequenced on an Illumina HiSeq 2000. We note that while all samples were run on the same flow cell, the samples were somewhat imbalanced by lane – however, the first PC of the expression data did separate perfectly by sampling location. Sample-specific information on reads and alignments are available in S1 Table.

### RNA-seq data generation

Resulting reads were mapped to the genome using TopHat2 (29) using the paired-ends procedure (we used the option --library-type fr-firststrand). Gene counts relative to the UCSC hg19 knownGene annotation were calculated using the featureCounts script of the subread package (version 1.4.6) (30). There were 23,710 genes in this annotation, and we dropped 305 genes that were annotated to more than 1 chromosome. Of the remaining 23,405 genes, 18,316 genes had non-zero expression counts in at least one sample. Counts were converted to FPKM (fragments per kilobase per million reads mapped) values to allow comparisons across genes with different lengths and libraries sequenced to different depths. These FPKMs were transformed prior to statistical analysis: *log*_2_(*FPKM* + 1). The log transformed FPKM values were used in all subsequent analyses.

### RNA-seq data analysis

Differential expression for sampling location was identified using Eq. 1 above, where *y*_*ij*_ represents transformed expression (rather than DNAm) levels, and different SVs (N=4) were calculated from the expression data. We carried out gene ontology analysis on the differentially expressed genes with the GOstats package (44). Transformed FPKMs were used to assess functional significance of differentially methylated features. We mapped the DMPs to genes in the UCSC knownGenes (hg19) and determined which DMPs exhibit correlation between DNAm and gene expression with the Matrix EQTL package (45). We used Pearson’s Chi-squared test with Yates’ continuity correction to examine whether DMPs are more likely to exhibit correlations between DNAm and gene expression than non-DMPs. We then mapped significant DMRs to genes expressed in the RNA-seq data (e.g. showing non-zero expression levels in ≥ 1 samples), and correlated the average DNAm level within the DMR to the transformed expression level. When multiple genes were within or near a DMR, we retained the gene (and its correlation) with the largest absolute correlation. We carried out gene ontology analysis for the genes proximal to DMRs with the GOstats package. For each significant block, we found the UCSC annotated gene(s) containing within the block and their evidence for differential expression as calculated above. We used Pearson’s Chi-squared test with Yates’ continuity correction to test whether differentially expressed genes were enriched in blocks compared to the rest of the transcriptome.

### Processing of public data and distance calculations

We performed a second larger data processing and normalization procedure on our scalp- and dura-derived fibroblasts after adding data from skin fibroblasts (GSE52025) (17), pure populations of blood (18) and prefrontal cortex cells (19) from the FlowSorted.Blood.450k and FlowSorted.DLPFC.450k Bioconductor packages respectively, and then melanoma data from TCGA (33). The M and U channels were combined across all experiments and then normalized with stratified quantile normalization as described above. We then dropped the probes on the sex chromosomes as well as probes that are common SNPs (based on dbSNP 142) as described above. Within the normalized data, we then calculated all pairwise Euclidean distances on the proportion methylation scale, and selected specific comparisons to display in Fig. 4.

### Differential variability and age related changes by tissue type

We calculated differential variability between scalp and dura CpG DNAm levels using the Levene test (46) and subsequent p-values were adjusted for multiple testing using the FDR. We filtered out the 101,989 genome-wide significant probes showing mean methylation differences by sampling location, as there is a strong mean-variation relationship in DNAm data due to being constrained within 0 and 1 (e.g. gaining methylation from an unmethylated state or losing methylation from a fully methylated state increases variance).

We tested for probes that showed differential age-related divergence in DNAm by fibroblast sampling location. First, we calculated the difference in DNAm between scalp- and dura-derived fibroblasts from the same individual at every probe (creating a 456,513 probe by 10 individual matrix). We then computed 3 surrogate variables (the number estimated by the SVA algorithm) for a statistical model with donor age, and fit the following linear model:

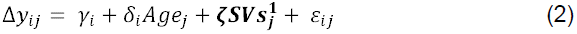

where Δ*y*_*ji*_ is the difference in DNAm levels between scalp and dura for probe *i* and individual *j*, *y*_*i*_ is the difference in DNAm levels at birth, *Age*_*i*_ is the age of the donor, and *δ*_*i*_ is the change in the difference of DNAm per year of life. We then generated a Wald statistic and corresponding p-value for *δ*_*i*_ and adjusted for multiple testing via the FDR. Post hoc age-related changes, e.g. the change in DNAm levels per year of life, were calculated within the scalp and dura samples. We then associated expression of nearby genes (within 5kb) with the DNAm levels at the probes showing significant age-by-location effects and performed gene ontology on the significant genes with the GOstats package (44).

### Variant calling

We called expression variants directly from the RNA sequencing alignments using samtools (version 1.1) and mpileup across all samples (47). We then filtered variants in the resulting variant call format (VCF) file based on coverage (<20), variant distance bias (p<0.05), read position bias (p<0.05), mapping quality bias (p<0.05), base quality bias (p<0.05), inbreeding coefficient binomial test (p<0.05), and homozygote bias (p>0.05). The resulting 64 high quality variants were annotated with SeattleSeq138 (38).

### Study approval

For every subject from whom the post-mortem tissues were collected, informed consent was obtained verbally from the legal next-of-kin using a telephone script, and was both witnessed and audiotaped, in accordance with the IRB approved NIMH protocol 90-M-0142 and the Department of Health and Human Services for the State of Maryland (protocol # 12-24).

### Data Availability

DNA methylation data in both raw and processed forms will be deposited on the Gene Expression Omnibus (GEO), accessing number pending. RNA sequencing reads will be deposited on the Short Read Archive (SRA, BioProject: PRJNA286856) and the genes counts will be deposited on GEO.

## Supporting Information

**S1 Fig. Experimental setup.**

We took dura and scalp samples from 11 donors ranging from 0.1 to 85 years of age. We then extracted and cultured fibroblasts from these samples, and performed genome-wide DNA methylation and RNA sequencing procedures on these fibroblasts.

**S2 Fig. DMR plots.**

DNA methylation levels (proportion methylation) of all 697 significant DMRs (FWER < 10%).

**S3 Fig. DNA methylation “blocks” plots.**

DNA methylation levels (proportion methylation) of all 243 significant differentially methylated blocks (FWER < 10%).

**S4 Fig. Principal Component Analysis Plots.**

The first principal component (PC1) of the gene expression data plotted against fibroblast sampling location (scalp versus dura). The first PC of the gene expression data mimics the first PC of the DNAm data; both represent sampling location.

**S5 Fig. Clustering analysis on DNAm data from cells of different tissues.**

(A) PC1 with respect to PC2 of the DNAm data from the following cells: various cells of the blood; neuronal (NeuN+) and glial (NeuN-) cells from the DLPFC; cultured fibroblasts derived from skin, dura mater, and scalp; cells from a primary solid skin tumor. (B) Cluster dendrogram constructed from the DNAm data from the cells in panel A.

**S6 Fig. Epigenomic distance within scalp-derived fibroblasts with respect to age differences between subjects.**

**S7 Fig. Age related DNAm divergence.**

DNAm plotted with respect to age for all 7,265 CpGs significantly associated with diverging DNAm levels across aging (at FDR < 10%).

**S8 Fig. Number of coding variants with respect to subject age.**

**S1 Table. Tissue donor demographics and RNAseq read alignment data.**

**S2 Table. Information on significant DMRs (FWER < 10%).**

**S3 Table. DMR Gene Ontology.**

Gene Ontology on genes that overlap or are proximal to (within 5kb) of significant DMRs (FWER < 10%).

**S4 Table. Gene Ontology on genes differentially expressed between scalp-and dura-derived fibroblasts (FDR < 5%).**

**S5 Table. DMR Gene Ontology.**

Gene Ontology on genes that overlap or are proximal to DMRs (within 5 kb) and exhibit significant correlation between gene expression and DNAm (p < 0.05).

**S6 Table. Gene Ontology on genes whose expression is correlated with nearby diverging DNAm CpGs.**

Gene Ontology on genes that overlap or are proximal to (within 5kbs) of CpGs that exhibit location-dependent age-related changes (FDR < 10%) and demonstrate correlation between DNAm and expression (p < 0.05).

**S7 Table. Candidate exonic variants between scalp- and dura-derived fibroblasts.**

